# Preliminary *in silico* experiments: Towards new cancer treatments?

**DOI:** 10.1101/2021.04.06.438636

**Authors:** Michel Fliess, Cédric Join, Kaouther Moussa, Seddik M. Djouadi, Mohamed Alsager

**Affiliations:** LIX (CNRS, UMR 7161), École polytechnique, 91128 Palaiseau, France; CRAN (CNRS, UMR 7039)), Université de Lorraine, BP 239, 54506 Vandœuvre-lès-Nancy, France; Université Grenoble Alpes, CNRS, Grenoble INP, GIPSA-lab, 38000 Grenoble, France; Department of Electrical Engineering and Computer Science, University of Tennessee, Knoxville, TN 37996, USA (,); AL.I.E.N., 7 rue Maurice Barrès, 54330 Vézelise, France (e-mail: {, })

**Keywords:** Cancer, chemotherapy, immunotherapy, flatness-based control, model-free control

## Abstract

We present some “in silico” experiments to design combined chemo- and immunotherapy treatment schedules. We introduce a new framework by combining flatness-based control along with model-free control. The flatness property of the used mathematical model yields straightforward reference trajectories. They provide us with the nominal open-loop control inputs. Closing the loop via model-free control allows to deal with the uncertainties on the injected drug doses. Several numerical simulations illustrating different case studies are displayed. We show in particular that the considered health indicators are driven to the safe region, even for critical initial conditions. Furthermore, in some specific cases there is no need to inject chemotherapeutic agents.

## I. Introduction

Mathematical modeling in oncology is a well established discipline (see, *e.g*., [Altrock *et al*. (2015)] for recent advances). We consider the problem of drug injections scheduling from a control point of view. Among the many models which have been used, those stemming from the earlier work of [Stepanova (1979)] are quite popular. Most appealing are several publications by d’Onofrio and different coauthors: see especially [d’Onofrio *et al*. (2012)]. Such approaches to chemo- and immunotherapy led in recent years to promising control-theoretic investigations: see, *e.g*., [Alamir (2014)], [Schättler & Ledzewicz (2015)] and references therein, [Moussa *et al*. (2020)], [Sharifi *et al*. (2017)], [Sharifi *et al*. (2020a)]. They employ various optimization techniques which are related to optimal control, model predictive control, and robust control.

We explore here another route via tools which are combined for the first time and do not seem to have been used before in oncology, although they gave rise to an abundant literature in control engineering:

1. *Flatness*-based control (see [Fliess *et al*. (1995)], [Fliess *et al*. (1999)], and [Lévine (2009)], [Sira-Ramírez & Agrawal (2004)]) has been well received in many industrial domains. See, *e.g*., [Bonnabel & Clayes (2020)] for tower cranes.
2. Besides being useful in concrete case-studies (see, *e.g*., [Amasyali *et al*. (2020)], [Park *et al*. (2021)], [Park & Olama (2021)] for energy management), *model-free* control in the sense of [Fliess & Join (2013)] has already been illustrated by drug injections for some type-1 diabetes [MohammadRidha *et al*. (2018)] and for inflammation [Bara *et al*. (2018)]. Note that the terminology “model-free control” has been used many times with different definitions: see, *e.g*., [Chareyron & Alamir (2009)] in oncology.

Our *virtual patient* is modeled through two ordinary differential equations presented in [d’Onofrio *et al*. (2012)]. This system is trivially flat with obvious *flat outputs*. The design of suitable reference trajectories with the corresponding open-loop controls becomes straightforward. A major source of uncertainty, according to [Sharifi *et al*. (2020a)], is the unknown fluctuation of the drug delivery to the tumor, which should be related to actuators faults, *i.e*., to a classic topic in fault-tolerant control (see, *e.g*., [Noura *et al*. (2015)]). It has been already noticed that model-free control is well-suited for dealing with actuators faults: see [Fliess & Join (2013)] for an academic example and [Lafont *et al*. (2015)] for a concrete case-study. The loop is therefore closed via model-free control. Let us emphasize the following points:

- the computer implementation is easy;
- only a low computing cost is necessary;
- some *scenarios, i.e., in silico* experiments, lead to unexpected results and should attract the attention of cancerologists. Our paper is organized as follows. Section II reviews briefly modeling, flatness-based control, and model-free control. Numerical simulations are presented in Section III. Section IV contains some suggestions for future research.

## II. Modeling and control

### A. Modeling for combined chemo- and immunotherapy

We consider the model presented in [d’Onofrio *et al*. (2012)]

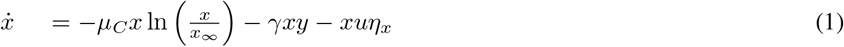

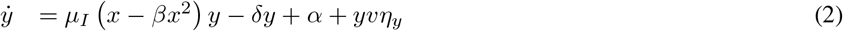

*x, y* are, respectively, the number of tumor cells and the immune cell density; the control variables *u* and *υ* are the cytotoxic and immune-stimulation drugs; the parameters *μ_C_, μ_I_, α, γ, δ, x*_∞_ are positive. The terms *η_x_, η_y_*, 0 ≤ *η_x_* ≤ 1, 0 ≤ *η_y_* ≤ 1, are inspired by [Sharifi *et al*. (2020a)]: they represent the uncertain and fluctuating parts of drugs which are delivered to the tumor.

### B. Flatness property

A control system with *m* independent control variables is said to be *(differentially) flat* if, and only if, there exists *m* system variables *y*_1_,…, *y_m_*, the *flat outputs*, such that any system variable *z*, the control variables for instance, may be expressed as a *differential* function of *y*_1_,…, *y_m_*, *i.e*., 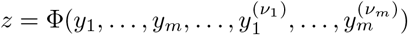, where the derivation orders *ν*_1_,…, *ν_m_* are finite. A linear system is flat if, and only if, it is controllable. Thus flatness may be viewed as another extension of Kalman’s controllability.

Equations (1)–(2) yield

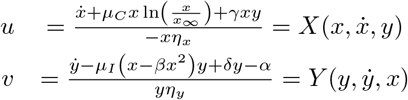

The above equations show immediately that System (1)-(2) is flat; *x, y* are flat outputs.

### C. Reference trajectory and nominal open-loop control

One of the main benefits of flatness is the possibility of easily deriving a suitable reference trajectory and the corresponding nominal open-loop control. For a given reference trajectory *x*⋆(*t*), *y*⋆(*t*), the corresponding nominal control variables *u*⋆(*t*) = *X*(*x*⋆(*t*), *ẋ*⋆(*t*), *y*⋆(*t*)), *υ*⋆(*t*) = *Y*(*y*⋆(*t*), *ẏ*⋆(*t*), *x*⋆(*t*)) might exhibit unacceptable negative values. Define therefore the nominal open-loop control variables *u*_OL_(*t*) = *u*⋆(*t*) if *u*⋆(*t*) ≥ 0, *u*_OL_(*t*) =0 if *u*⋆(*t*) < 0, and *υ*_OL_(*t*) = *υ* ⋆ (*t*) if *υ*⋆(*t*) ≥ 0, *υ*_OL_(*t*) = 0 if *υ*⋆(*t*) < 0.

### D. Closing the loop via model-free control

From a control-engineering standpoint the terms *η_x_* and *η_y_* should be related to actuators faults. Introduce therefore the two “decoupled” *ultra-local models* ([Fliess & Join (2013)], [Lafont *et al*. (2015)]):

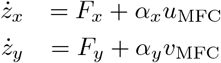

where *z_x_* = *x* – *x*⋆, *z_y_* = *y* – *y*⋆ are the tracking errors; *α_x_* (resp. *α_y_*) is a constant parameter which is chosen by the practitioner such that *ẋ* and *α_x_u* (resp. *ẏ* and *α_y_υ*) are of the same order of magnitude; *F_x_* and *F_y_*, which are data-driven, subsume the poorly known structures and disturbances. A real-time estimation ([Fliess & Join (2013)]) of *F_x_, F_y_* are given by

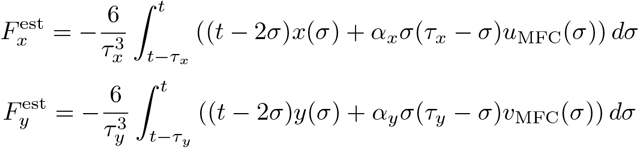

where *τ_x_, τ_y_* > 0 are “small.” Close the loop via an *intelligent Proportional* controller, or *iP*,

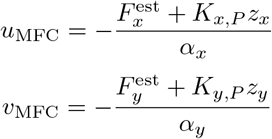

where *K_x, P_, K_y, P_* > 0. From

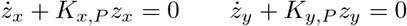

it follows that those two gains ensure local stability around the reference trajectory.

The close-loop controls *u*_CL_, *υ*_CL_ may now be defined:

- if *u*_OL_ + *u*_MFC_ ≥ 0, then *u_CL_* = *u*_OL_ + *u*_MFC_;
- if *u*_OL_ + *u*_MFC_ < 0, then *u*_CL_ = 0;
- if *υ*_OL_ + *υ*_MFC_ ≥ 0, then *υ*_CL_ = *υ*_OL_ + *υ*_MFC_;
- if *υ*_OL_ + *υ*_MFC_ < 0, then *υ*_CL_ = 0.

## III. Numerical simulations

### A. Presentation

A huge number in silico experiments have been most easily performed. Only a few of them are presented below. Like in several other papers, the parameters in System (1)-(2) are given in Table 1 ([d’Onofrio *et al*. (2012)]). There are three equilibria corresponding to *ẋ* = *ẏ* = *u* = *υ* = 0: 1)a locally stable equilibrium *x* = 73, *y* = 1.32 which corresponds to a benign case; 2) an unstable saddle point *x* = 356.2, *y* = 0.439, which separates the benign and malignant regions; 3) a locally stable equilibrium *x* = 737.3, *y* = 0.032, which is malignant. The duration of an experiment is 60 days. The time sampling interval is equal to 30 minutes.

**TABLE I:**
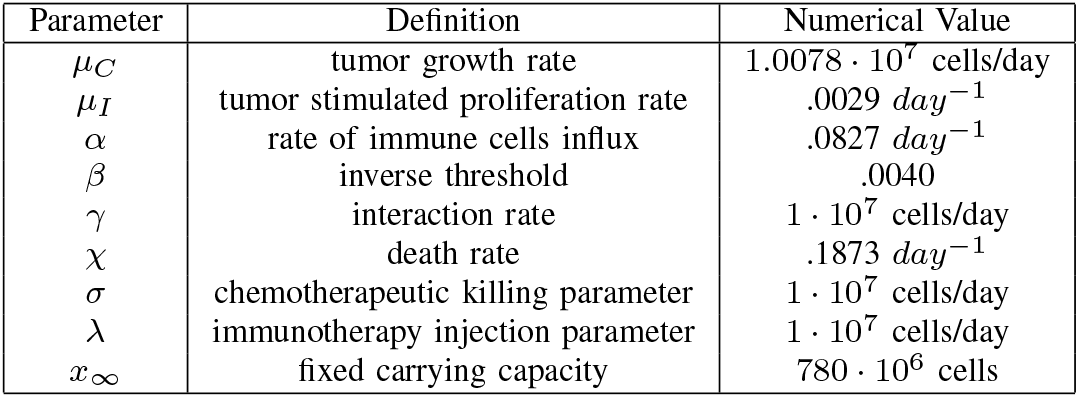

| Parameter | Definition | Numerical Value |
| --- | --- | --- |
| $\mu_C$ | tumor growth rate | $1.0078 \cdot 10^7$ cells/day |
| $\mu_I$ | tumor stimulated proliferation rate | $.0029 \text{ day}^{-1}$ |
| $\alpha$ | rate of immune cells influx | $.0827 \text{ day}^{-1}$ |
| $\beta$ | inverse threshold | .0040 |
| $\gamma$ | interaction rate | $1 \cdot 10^7$ cells/day |
| $\chi$ | death rate | $.1873 \text{ day}^{-1}$ |
| $\sigma$ | chemotherapeutic killing parameter | $1 \cdot 10^7$ cells/day |
| $\lambda$ | immunotherapy injection parameter | $1 \cdot 10^7$ cells/day |
| $x_\infty$ | fixed carrying capacity | $780 \cdot 10^6$ cells |

### B. Closed-loop and total amount of drugs

Set *η_x_* = *η_y_* = 0.5. This nominal value might be large according to [Sharifi *et al*. (2020a)]. Figures 1 and 2 display two experiments with the same initial point *x* = 500, *y* = 0.5, which lies in the attraction region of the malignant equilibrium. The total amounts of injected drugs, which are often considered as important constraints, are given by the two integrals 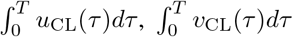, where *T* is the experiment duration. Figure 3 indicates that the quantity of drugs injected during the slow scenario is lower than in the fast one. This outcome ought to be discussed in oncology.

**Fig. 1:**
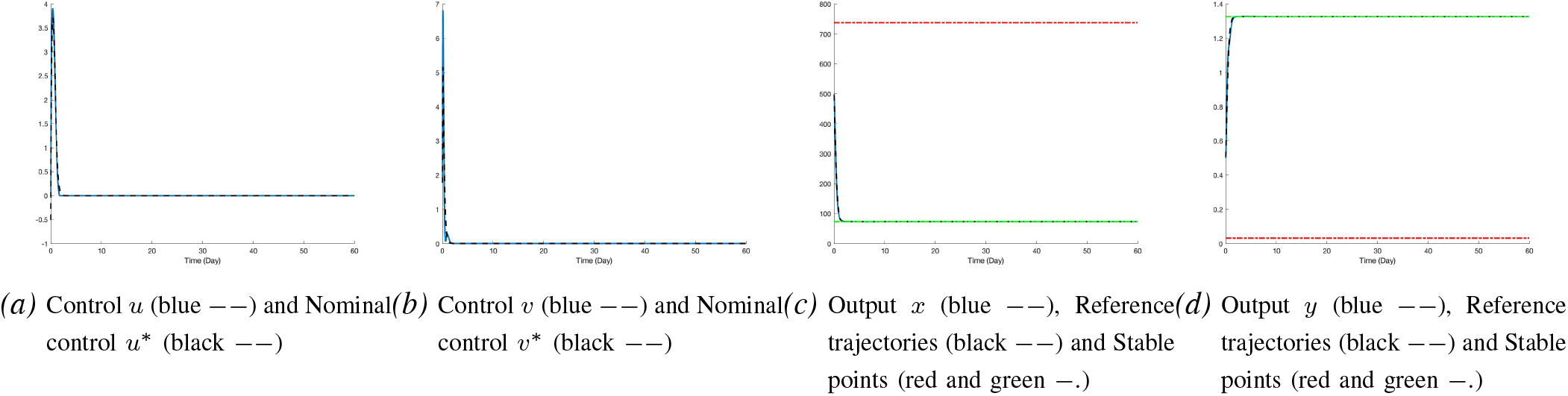
Fast trajectory

**Fig. 2:**
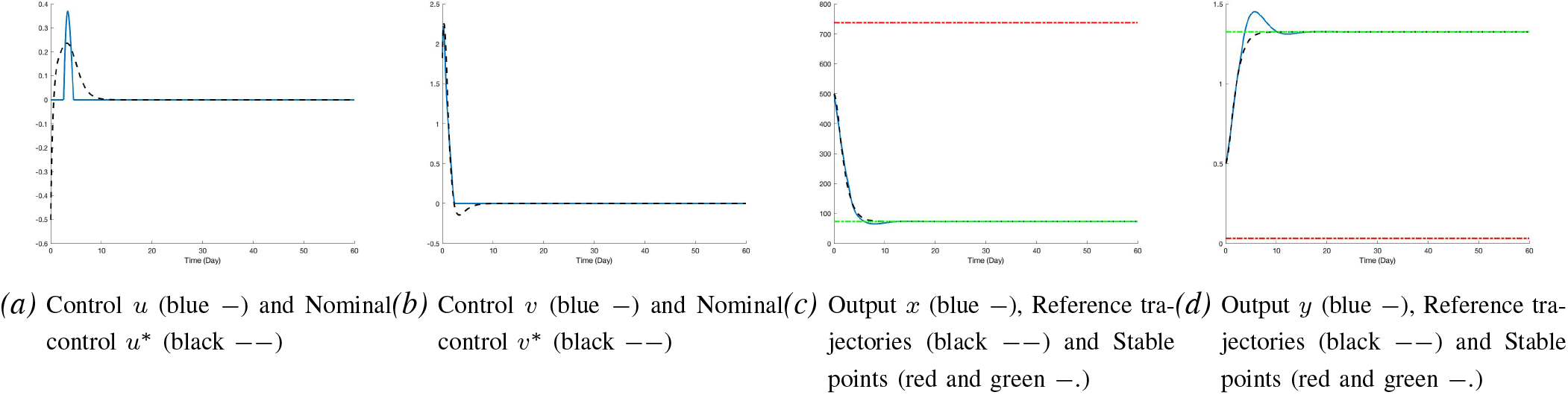
Slow trajectory

**Fig. 3:**
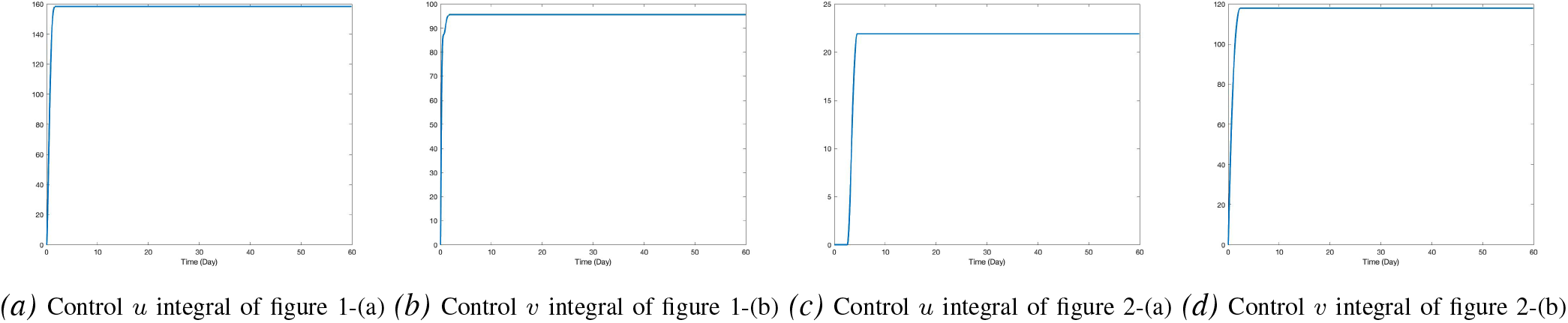
Comparison between total drug injections

### C. Other scenarios

1. *Same initial point.:* Here *η_x_* = 0.25, *η_y_* = 0.75 are supposed to be unknown. Use the same nominal parameters as in Section III-B, and the feedback loop of Section II-D, with *α_x_* = −10000, *α_y_* = 1, *K_x,P_* = 100, *K_y,P_* = 10. The results depicted in Figures 4 and 5 show that the benign equilibrium is reached after a short period of time.
2. *New initial point.:* The virtual patient is in a critical state, *i.e*., the initial state *x* = 770, *y* = 0.1 is close to the malignant equilibrium. The time variation of *η_x_* and *η_y_*, which are displayed in Figure 8, are assumed to be unknown. It is possible to cure the virtual patient without the cytotoxic drug, *i.e*., *u*_CL_ ≡ 0. Figure 6, which should be of interest for cancerologists, exhibits a convergence to the benign equilibrium with some oscillations perhaps due to the violent fluctuations of *η_y_*. The quality of the open loop behavior in Figure 7 is lower.

**Fig. 4:**
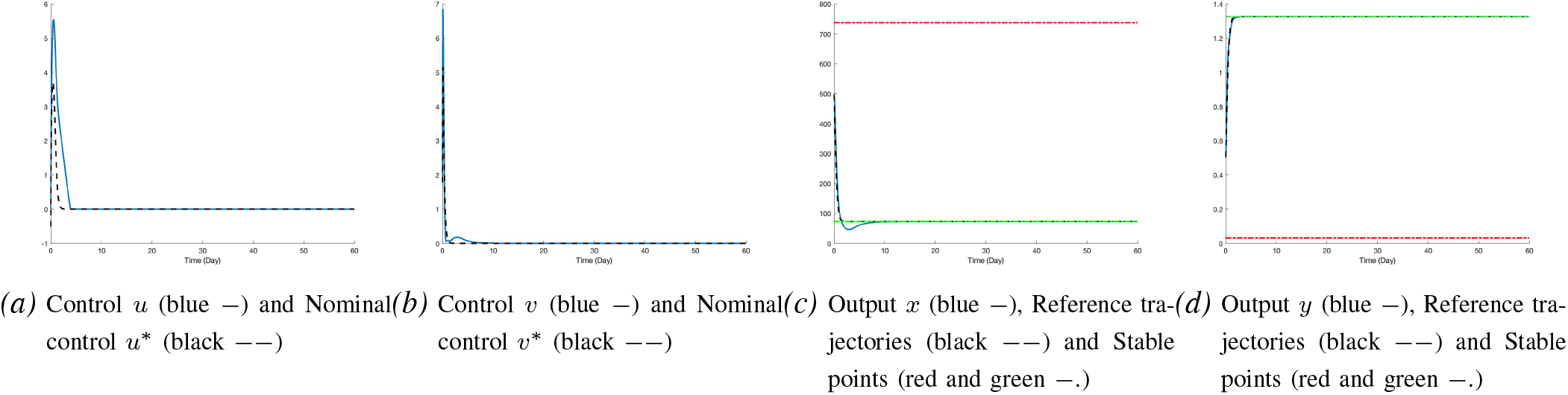
Unknown variation of *η_x_*

**Fig. 5:**
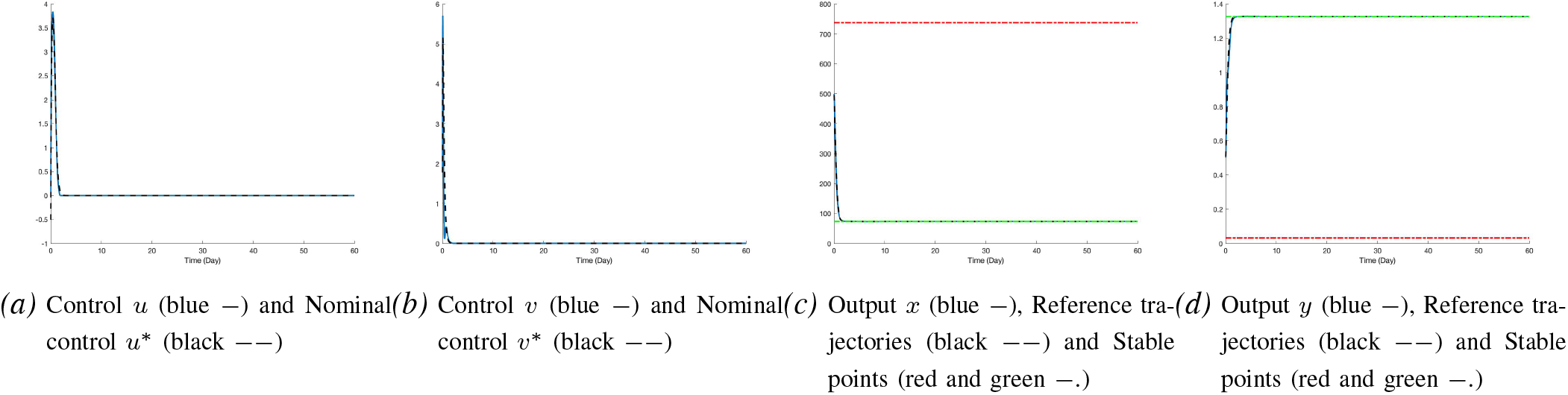
Unknown variation of *η_y_*

**Fig. 6:**
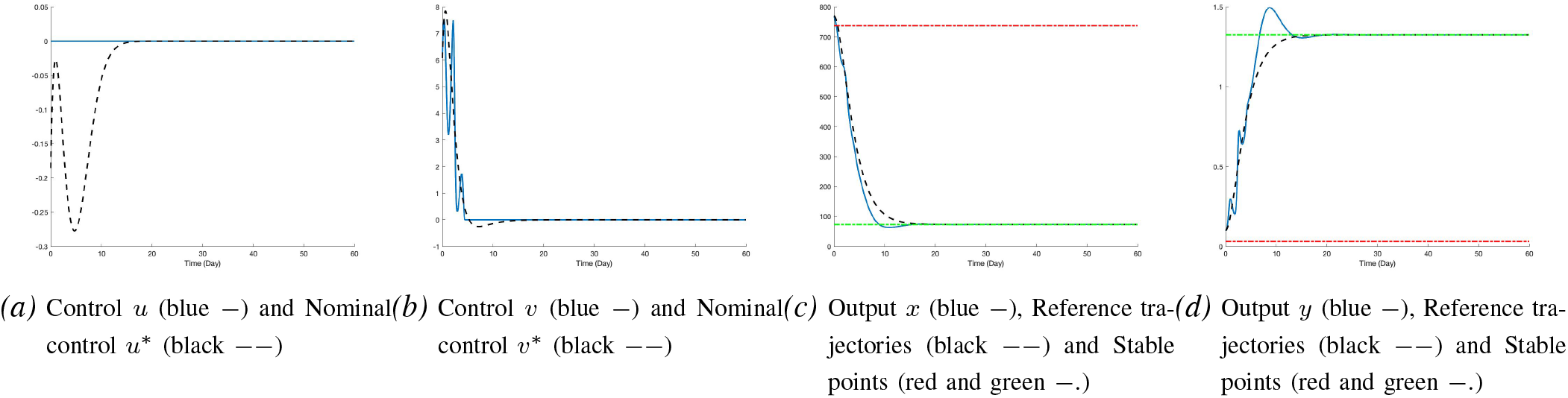
Very sick patient

**Fig. 7:**
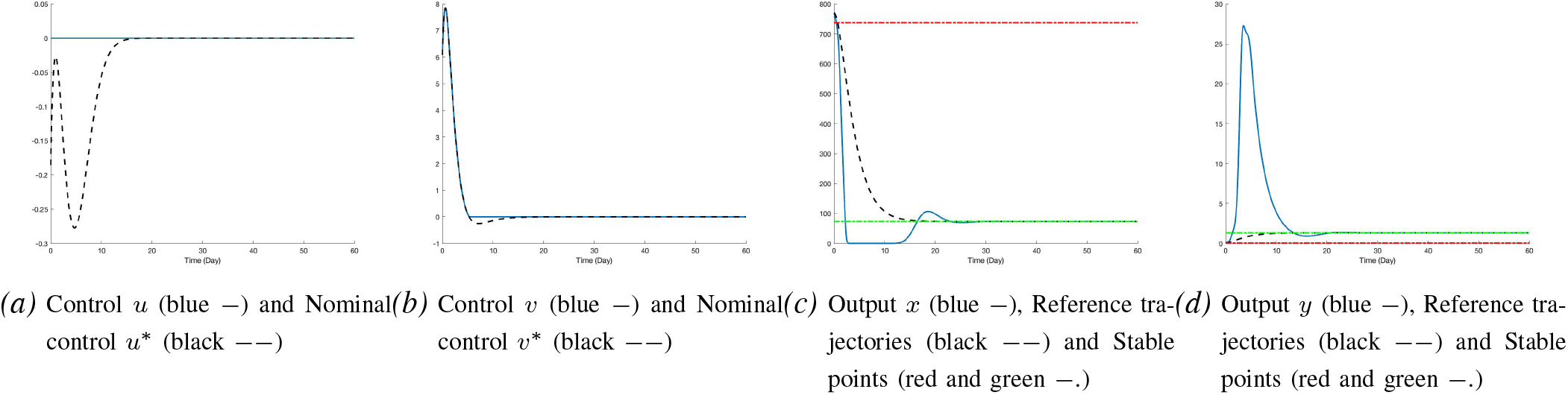
Open loop

**Fig. 8:**
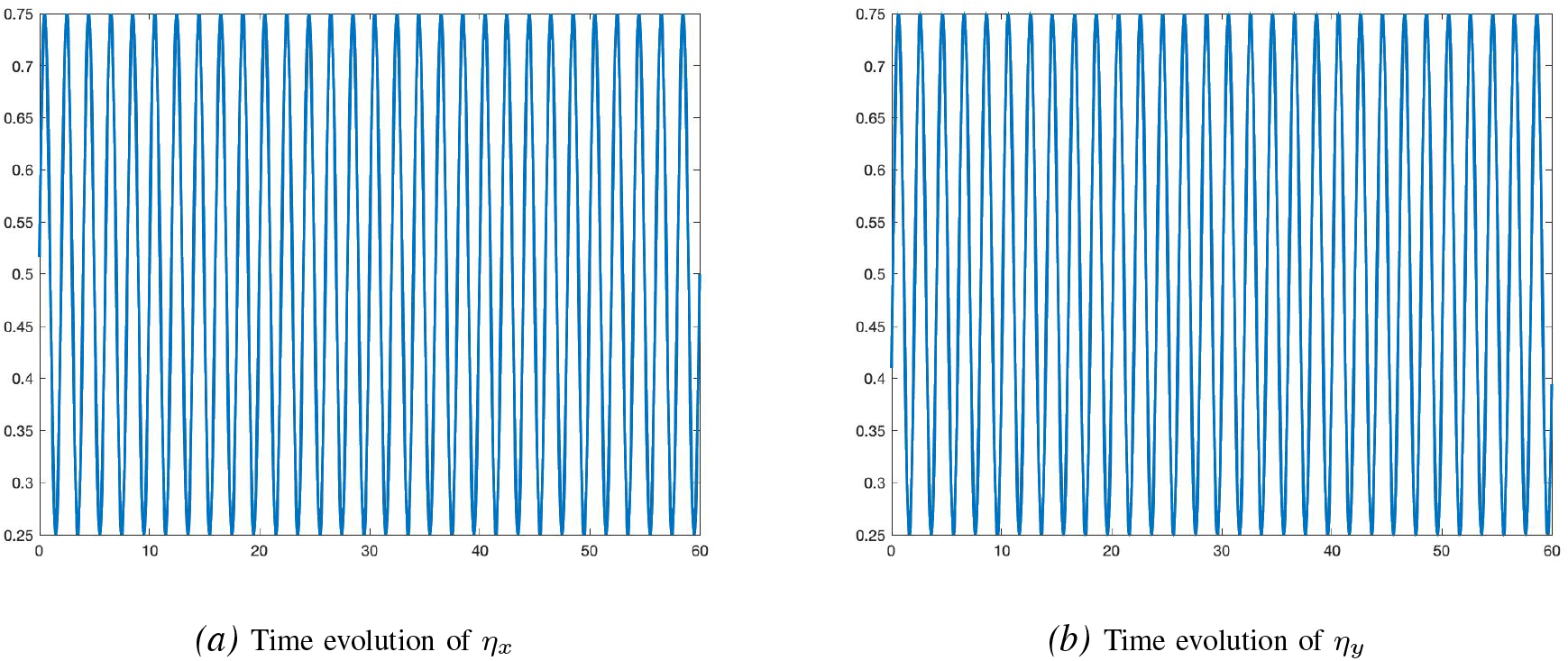
Fluctuation of the drug delivery

## IV. Conclusion

### A. Main goal

According to our computer experiments, there are critical situations where only immunotherapy matters: the cytotoxic drugs are useless. Those startling calculations need of course to be further analyzed.

### B. Systems Biology

In the spirit of *Systems Biology* (see, *e.g*., [Del Vecchio & Murray (2015)]), let us suggest the the following research tracks:

- Examine the parameter identification in Equation (1).
- Lack of time prevented us for examining more thoroughly the unavoidable constraints especially for the cytotoxic and immune-stimulation drugs. It will be done in a future publication thanks to the optimization techniques associated to flatness-based control (see, *e.g*., [Petit *et al*. (2001)], [Sira-Ramírez & Agrawal (2004)]).
- Flatness-based control might be helpful elsewhere:

1. Another model due to [Hahnfeldt *et al*. (1999)] has also been investigated from a control-theoretic perspective (see, *e.g*., [Kovács *et al*. (2014)], [Schättler & Ledzewicz (2015)] and references therein, [Cacace *et al*. (2018)]). It is easy to check that it is flat.
2. The unicycle in [Sharifi *et al*. (2020b)], which is used as a nanorobot for drug delivery, is well known to be flat.

